# Revisiting the intrageneric structure of the genus *Pseudomonas* with complete whole genome sequence information: Insights into Diversity and Host-related Genetic Determinants

**DOI:** 10.1101/2020.06.26.172809

**Authors:** Buqing Yi, Alexander H. Dalpke

**Author notes:** Correspondence: Dr Buqing Yi, Institute of Medical Microbiology and Virology, Medical Faculty, Technical University Dresden, Dresden, Germany. Fetscherstraße 74, 01307 Dresden, Germany.

## Abstract

*Pseudomonas* spp. exhibit considerable differences in host specificity and virulence. Most *Pseudomonas* species were isolated exclusively from environmental sources, ranging from soil to plants, but some *Pseudomonas* species have been detected from versatile sources, including both human host and environmental sources. Understanding genome variations that generate the tremendous diversity in *Pseudomonas* biology is important in controlling the incidence of infections. With a data set of 704 *Pseudomonas* complete whole genome sequences representing 186 species, *Pseudomonas* intrageneric structure was investigated by hierarchical clustering based on average nucleotide identity, and by phylogeny analysis based on concatenated core-gene alignment. Further comparative functional analyses indicated that *Pseudomonas* species only living in natural habitats lack multiple functions that are important in the regulation of bacterial pathogenesis, indicating the possession of these functions might be characteristic of *Pseudomonas* human pathogens. Moreover, we have performed pangenome based homogeneity analyses, and detected genes with conserved structures but diversified functions across the *Pseudomonas* genomes, suggesting these genes play a role in driving diversity. In summary, this study provided insights into the dynamics of genome diversity and host-related genetic determinants in *Pseudomonas*, which might help the development of more targeted antibiotics for the treatment of *Pseudomonas* infections.

## 1. Introduction

The gram-negative genus *Pseudomonas* (phylum Proteobacteria) constitutes a highly diverse group which is widely distributed in nature, utilizing a wide range of organic compounds as energy sources (1,2) and producing a large number of secondary metabolites (3).

*Pseudomonas* spp. are responsible for many plant diseases, but some *Pseudomonas* spp. may actually promote plant growth or protect plant from infections (4,5). A variety of *Pseudomonas* spp. may act as opportunistic human pathogen. For example, *P. putida* strains are prevalent in natural habitats, including soil, water, and plant surfaces, but have also been reported as opportunistic human pathogens causing nosocomial infections (6). *P. stutzeri* is a nonfluorescent denitrifying bacterium widely distributed in the environment, and it has been isolated as opportunistic pathogen from humans as well (7). Similarly, in addition to being frequently detected from many different environmental sources, *P. aeruginosa* acts as an important opportunistic human pathogen implicated in the pathogenesis of many diseases, e.g. in lung infections in cystic fibrosis (CF). In CF patients, dominance of *P. aeruginosa* in the lung microbial community correlates with multiple CF disease features, such as cough scores, chest radiograph scores, and loss of pulmonary function (8–10). Importantly, most *Pseudomonas* species that may act as human pathogens are resistant to multiple antibiotics, which makes it difficult to get rid of the infections they have caused, and controlling chronic *P. aeruginosa* airway infection in CF patients is still one of the most challenging tasks for CF treatment (11,12). It is in urgent need of looking for targets for developing effective antibiotics.

*Pseudomonas* can be transmitted via several distinct routes from animals, plants or environmental sources to human hosts. It is therefore important to understand the genome variations behind the highly diversified host options. Several previous studies have investigated *Pseudomonas* genome variations with different approaches, such as phylogeny based on multilocus sequence analysis (MLSA), average nucleotide identity (ANI), or average amino acid identity (13–21). These studies provided valuable information for *Pseudomonas* phylogeny research. However, the results of these studies are not completely consistent, which might be due to the differences in genome datasets and methods applied for phylogeny analysis among these studies. Owing to lack of complete whole-genome sequences when these studies were performed, only a few studies used whole-genome sequences for the genome variation analysis, and the assembly levels of most genomes included were scaffold. This means there is missing of genome information in certain regions, which could impact the results of genomic analysis. In this study, we aimed to investigate the intrageneric structure of *Pseudomonas* with a data set of 704 *Pseudomonas* complete whole genome sequences representing 186 species. With many whole-genome sequences available for *Pseudomonas* species with well-documented ecological information, we assume that genomic structure analysis might be able to provide insights into host-related genetic elements.

## 2. Materials and Methods

### Dataset

The analyzed data set consisted of 704 genomes of *Pseudomonas* strains available in January 2020 in the RefSeq database of the National Center for Biotechnology Information (NCBI). Only genomes with the assembly level of “complete genome” or “chromosome” are included in this data set. Accession numbers and other detailed information about the genomes are shown in Supplementary Table S1.

### Pangenome analysis

Pangenome analysis of *Pseudomonas* was performed with anvi’o (version 6.1) (22) following the workflow for microbial pangenomics (23,24). In detail, PGAP (25) was used to identify open reading frames, then each database was populated with profile hidden Markov models (HMMs) by comparing to a collection of single-copy genes using HMMER (26). We used the Diamond software (27) to calculate gene similarity and MCL (28) for clustering under the following settings: minbit, 0.5; mcl inflation, 10; and minimum occurrence, 2.

### Intrageneric structure analysis

Whole-genome comparisons by ANI (average nucleotide identity) was calculated for blast based ANI (ANIb) and MUMmer based ANI (ANIm) with PYANI (29). For phylogenomic analysis of *Pseudomonas* genomes, we used the sequence information of single-copy core genes present in 100% *Pseudomonas* stains. The program MUSCLE (30) with default settings was used for creating alignment of protein sequences. The alignments were cleaned up by removing positions that were gap characters in more than 50% of the sequences using trimAI (31). Sequence alignments of core genes were concatenated to give a single core alignment and a maximum likelihood phylogeny was then generated using the program IQ-TREE (32) with a time-reversible amino-acid substitution model WAG, four gamma categories for rate heterogeneity and 1000 bootstrap replicates. The WAG model was chosen because it has been shown to be applicable to phylogenetic studies of a broad range of protein sequences and generate more accurate phylogenetic tree estimates compared to many other models (33). All phylogenies were visualized using the Interactive Tree of Life (34).

### Comparative functional enrichment analysis

All genomes were first annotated with the functional annotation source COGs (35). Then functional enrichment analysis was performed using a Generalized Linear Model with the logit linkage function and Rao score test to compute an enrichment score and p-value for each function, then R package qvalue is used to do False Detection Rate correction to p-values to account for multiple tests.

### Functional and geometric homogeneity analyses

Both functional and geometric homogeneity analyses were carried out using functions for homogeneity computing from anvi’o (version 6.1), which are integrated in the workflow for microbial pangenomics (22,23). Functional homogeneity index indicates how aligned amino acid residues are conserved across genes. To perform this analysis, first, based on amino acids’ biochemical properties, all amino acids are divided into seven different conserved groups: nonpolar, aromatic, polar uncharged, both polar and nonpolar characteristics, acidic, basic, and mostly nonpolar. Next, based on how close the biochemical properties of the amino acid residues are to each other, for every pair of amino acids at the same position in the orthologous gene, a similarity score is assigned. Functional homogeneity index is then calculated with all assigned similarity scores and will reach its maximum value of 1.0 if all residues are identical. Geometric homogeneity index indicates how gene structure is conserved within one orthologous gene family. It is calculated based on the gap/residue distribution patterns, with a maximum value of 1.0 indicating there are no gaps in the alignment.

## Results and Discussion

### 1. Pan-genome characteristics of *Pseudomonas*

To investigate the genomic diversity of *Pseudomonas* spp., we compiled a total of 704 complete whole genomes (with assembly level “complete genome” or “chromosome”) downloaded from NCBI (see Table S1 in the supplemental material). This dataset contains genomes representing 186 species, among which 95 species have been validly published (verified through The Bacterial Diversity Metadatabase https://bacdive.dsmz.de/) (see Table S2 for a detailed species information). Across the entire data set, genome size varied between 3.19 and 7.76 Mb (mean, 6.32 Mb), GC content varied between 0.38 and 0.68 (mean, 0.63), the number of predicted genes ranged from 2,831 to 7,360 (mean, 5,637).

We performed pangenome analysis of the *Pseudomonas* dataset with anvi’o (version 6.1) (22) following the workflow for microbial pangenomics. We first analyzed the number of orthologous gene families that are shared by all the strains, and the results showed that only 122 genes are present in 100% *Pseudomonas* stains, which represents 0.2% of the entire pan-genome (62,202 genes). Based on a widely used criterion (36–38), orthologous gene families have been classified into core genome (present in 99% ≤ strains ≤100%) and accessory genome (present in < 99% stains). The core genome at the genus level is composed of 1009 genes (see Table S3 for detailed information), representing 1.62 % of the entire pan-genome. The large accessory genome (98.38% of the pan-genome) indicates a high level of genomic diversity of *Pseudomonas* genomes, which is consistent with the observed biological diversity of the *Pseudomonas*, highlighting the heterogeneity of this group of bacteria.

### 2. Intrageneric structure based on whole-genome sequences

To determine the degree of genomic relatedness and hence clarify the relationship between the *Pseudomonas* spp., we calculated the pairwise average nucleotide identity (ANI) for all possible pairs of genomes. ANI estimates the average nucleotide identity of all orthologous genes shared between any two genomes. This index is a robust similarity metric that has been broadly used to delineate inter- and intra-species strain relatedness (39).

We included all validly published *Pseudomonas* species in our dataset for ANI analysis. To achieve a better resolution of the intrageneric structure at the genus level, for species with multiple genomes, maximal three genomes from one species were randomly chosen and included in the ANI analysis, with the exception of all reference or representative genomes being included. Thus, we have performed ANI analysis with 155 representative strains from all 95 valid species in this dataset including 25 reference or representative genomes (see Table S4 for detailed information). Based on whole-genome similarity index ANIb, *Pseudomonas* strains clearly fall into four clades. Representative species in each clade are labelled in Figure 1. Likewise, the intrageneric structure derived by ANIm analysis comprises four clades (Figure S1), and the species fall into each clade are almost identical between ANIb and ANIm based analyses. Detailed information about clade assignment of each genome has been labelled in Table S4. Based on in-depth literature research about ecological information of each species, most species that have documented evidence for being identified as human or animal pathogens appear closely related and fall into clade1(as shown in Table 1, please see supplementary document for detailed literature information), such as *P. aeruginosa, P. stutzeri and P. putida*. Species falling into clade2 are from mixed environmental sources, ranging from soil to plants. Species in clade3 are mostly plant-associated bacteria, with functional feature as plant growth-promoting bacterium or plant-pathogen. Species in clade4 are mostly plant pathogens, such as *P. syringae* and *P. cerasi*.

**Fig. 1.**
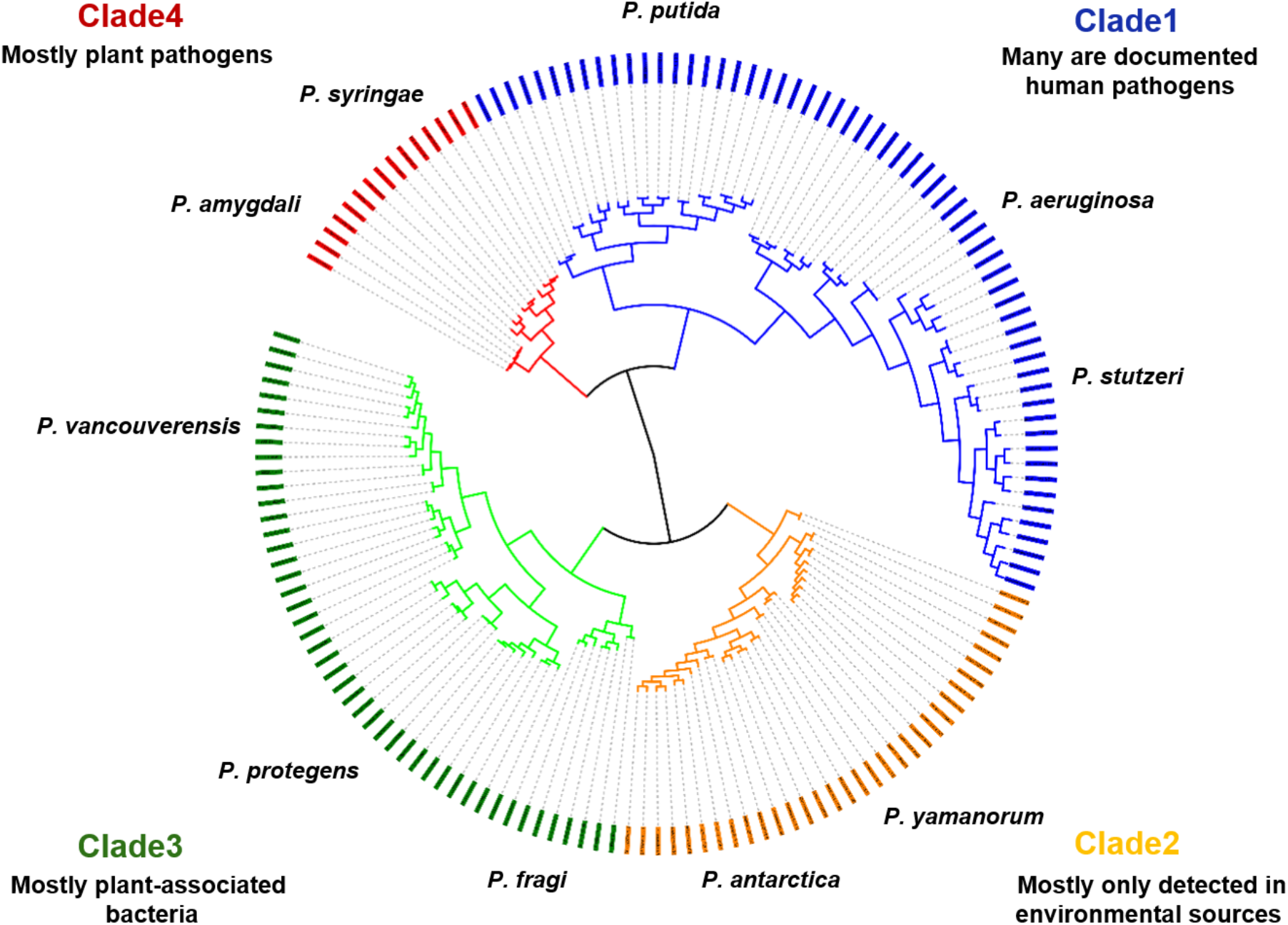
Intrageneric structure of the genus *Pseudomonas* based on whole-genome comparisons (ANIb) of 155 *Pseudomonas* strains representing 95 valid *Pseudomonas* species. The calculations were performed using the Python module PYANI. The strains fall into four major Clades (Clade1-Clade4). The positions of representative species in each Clade were labelled.

**Table 1.** Clade information summary.

|  | Number of genomes | Number of species | Number of species that may act as human or animal pathogen* |
| --- | --- | --- | --- |
| Clade1 | 61 | 39 | 18 |
| Clade2 | 33 | 19 | 3 |
| Clade3 | 46 | 28 | 3 |
| Clade4 | 15 | 9 | 0 |

\*Please see supplementary document for detailed literature information.

We also analyzed the intrageneric structure of the genus *Pseudomonas* using concatenated alignment of all single-copy core genes based on pan-genome analysis. As shown in Figure 2, phylogeny analysis of the 155 genomes also revealed four clades showing similar topology as ANI dendrograms (Figure 1 & Figure S1), with clade1 & clade4 more closely related, and clade2 & clade3 more closely related. The species falling into each clade are the same as that revealed by ANI analyses. Furthermore, we have constructed a *Pseudomonas* phylogeny tree with all the 704 genomes included in our dataset. Similarly, four clades can be identified based on the relative position of representative species (Figure S2).

**Fig. 2.**
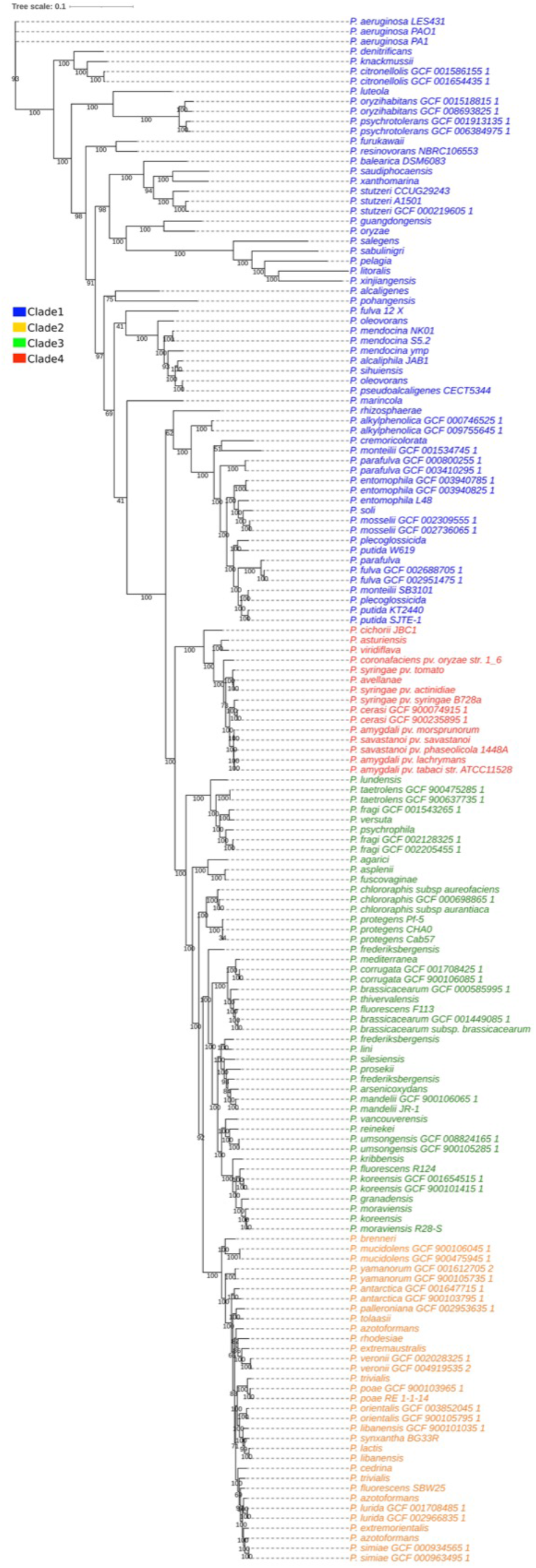
Phylogeny of 155 *Pseudomonas* genomes. The phylogenetic analysis was performed using the Maximum Likelihood method based on concatenated single-copy core genes and presented as an unrooted tree. Four clades comparable to the four major clades formed by whole-genome comparisons (ANI) have been detected and colorlabelled. The numbers indicate bootstrap values of the branches.

This intrageneric structure is congruent with those from several previously performed *Pseudomonas* phylogeny analyses in which a large number of genomes from NCBI were evaluated by either four concatenated genes (16S rRNA, gyrB, rpoB, and rpoD) or MLSA or average amino acid identity (14,17,21). In these studies, *Pseudomonas* species that may act as opportunistic human pathogens or plant pathogens are mostly separated from other nonpathogenic *Pseudomonas* species. Furthermore, in the current study, the intrageneric structure based on *Pseudomonas* phylogeny established with concatenated core-gene alignment is largely consistent with the intrageneric structure based on ANI. This is in line with one recent investigation about *Pseudomonas* phylogeny (14) reporting that ANI results show very similar topology as phylogeny analysis based on MLSA.

### 3. Comparative functional enrichment analysis reveals host-specific features

Among the *Pseudomonas* spp., most species were purely isolated from environmental sources ranging from soil to plants, but some species may act as opportunistic human pathogens causing infections in patients. It remains unclear why some species may infect humans or animals, but some other species are only able to live in natural habitats.

ANI based genome intrageneric structure analysis indicates that most species that may act as opportunistic human pathogen or use animals as host are closely related and fall into clade1 (Figure 1 & Figure S1). We asked whether there are any functions predominantly present in genomes from clade 1, while absent from other clades. Our hypothesis is these genes are host-related genetic elements playing an important role in the pathogenicity of *Pseudomonas* human pathogens. We therefore conducted comparative functional enrichment analyses. Because the numbers of genomes in each clade were greatly dissimilar, to avoid statistical bias, we first restricted our comparative analyses to five representative genomes from each clade, including all known reference genomes or representative genomes. The detailed information of these genomes is listed in Table S5. Their hosts or environments are representative of the diversity of ecological niches of *Pseudomonas*.

Comparative functional enrichment analyses showed that multiple genes are highly enriched in genomes from clade1, while predominantly absent in genomes from other clades consisting of species mainly living in natural habitats (Table S6). The results were verified with the 155 genomes for ANI analysis representing 95 valid *Pseudomonas* species, and all the genes were also highly enriched in genomes from clade1 in this dataset, while predominantly missing in genomes from other clades (Table S7). Since many *Pseudomonas* species have multiple strains, it is important to know if the same results can be achieved if more strains are used for the comparative functional enrichment analysis. We therefore prepared a dataset of 226 genomes from several species with multiple strains, including 113 randomly chosen genomes from clade1 (36 *P. putida* genomes, 18 *P. stutzeri*, 6 *P. monteilii* and 53 *P. aeruginosa*) and 113 genomes from clade 2/3/4 (10 *P. synxantha* genomes, 39 *P. syringae*, 6 *P. orientalis*, 51 *P. chlororaphis* and 7 *P. brassicacearum*). Equal number of genomes from clade1 and clade2/3/4 were included in this dataset to avoid statistical bias (see Table S8 for the information of these genomes). 95% of the functions that are predominantly enriched in human pathogens have been further verified with this multi-strain dataset (details shown in Table S9). For example, all the subunits of the Na+ transporting NADH:ubiquinone oxidoreductase (nqrA/nqrB/nqrC/nqrD/nqrE/nqrF) are present in all *P. aeruginosa* and *P. stutzeri* strains, while missing in all the strains from clade2/3/4. This result is consistent with what we know from *P. aeruginosa* that there is little association between a strain’s genotype and the niche of its isolation source, indicating that the ability of *P. aeruginosa* to infect humans is a feature of *P. aeruginosa* as a species, not a characteristic of specific *P. aeruginosa* strains (40,41).

The most consistent results from the three analyses with 20 representative genomes, 155 genomes for ANI analysis and 226 genomes with multiple strains are shown in Figure 3. These genes include: ybcL (protein: Membrane-bound metal-dependent hydrolase Ybcl); nqrA/nqrB/nqrC/nqrD/nqrE/nqrF (Na+-transporting NADH:ubiquinone oxidoreductase (NQR), subunit NqrA/NqrB/NqrC/NqrD/NqrE/NqrF); proQ (ProQ activator of osmoprotectant transporter, sRNA-binding protein); rhlB (UDP:flavonoid glycosyltransferase YjiC); crfC (Replication fork clamp-binding protein CrfC, dynamin-like GTPase family); rbn/rnz/elaC (Ribonuclease BN, tRNA processing enzyme); MFS_1 homolog (ATP/ADP translocase/MFS transporter), and two uncharacterized conserved genes ykaA and ydbL. Here gene names refer to representative homolog names in *E. coli* or in one *Pseudomonas* species.

**Fig. 3.**
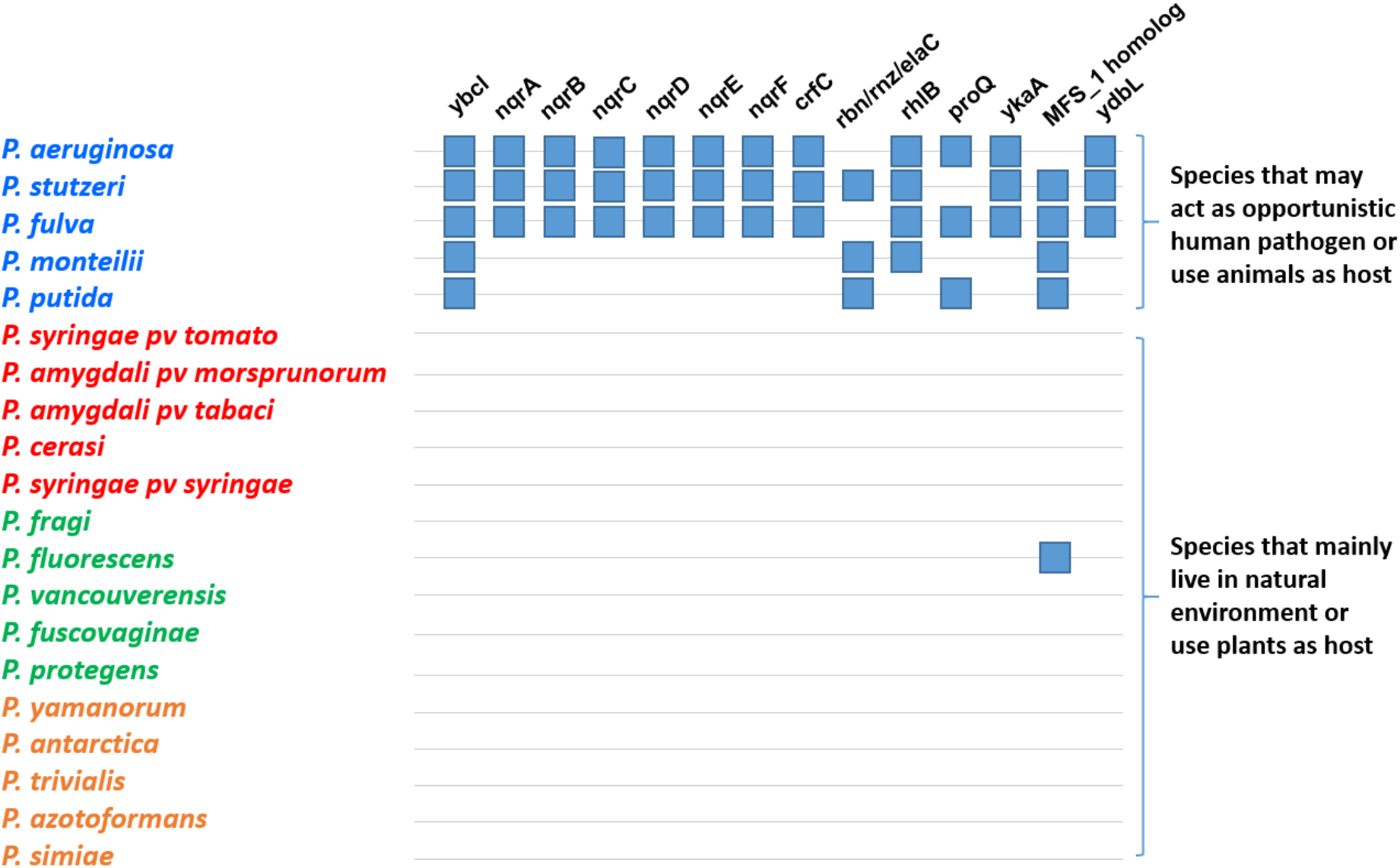
Distribution of differentially enriched functions in 20 representative *Pseudomonas* species. Species colors are indicative of the ANI based clade that the respective species fall into, with clade 1 labelled as blue, clade 2 as yellow, clade 3 as green and clade4 as red. Squares correspond to functions that were found to be preferably enriched in species that may act as opportunistic human pathogen or use animals as host. From left to right: ybcl (Membrane-bound metal-dependent hydrolase Ybcl); nqrA/nqrB/nqrC/nqrD/nqrE/nqrF (Na+-transporting NADH:ubiquinone oxidoreductase (NQR), subunit NqrA/NqrB/NqrC/NqrD/NqrE/NqrF); crfC (Replication fork clamp-binding protein CrfC, dynamin-like GTPase family); rbn/rnz/elaC (Ribonuclease BN, tRNA processing enzyme); rhlB (UDP:flavonoid glycosyltransferase YjiC); proQ (ProQ activator of osmoprotectant transporter, sRNA-binding protein); ykaA (uncharacterized conserved protein YkaA); MFS_1 homolog (ATP/ADP translocase/MFS transporter); ybdL (uncharacterized conserved protein YdbL).

Consistent with our hypothesis, most of these genes are involved in the regulation of bacterial pathogenesis, not only in *Pseudomonas* pathogens, also in other well-known human pathogens. Among these genes, rhlB and nqrB are proved virulence factors in *P. aeruginosa* (42). Furthermore, it has been recently reported that *P. aeruginosa* NADH:ubiquinone oxidoreductase (NQR) is actually a completely new form of proton pump, which plays an important role in the adaptation against autotoxicity during the infection process (43). ProQ is a major small RNA-binding protein in bacteria (44). It has been shown to target hundreds of transcripts, including mRNAs from many virulence regions in various pathogenic bacteria (45– 48). In *Salmonella* infected human cells, ProQ not only regulates bacterial motility, chemotaxis, and virulence gene expression, also affects MAPK signaling pathways in the host, suggesting a critical role of ProQ in regulating pathogenesis (49). In line with the role of small RNA-binding protein ProQ, it has been shown that the major function of ribonuclease BN is to control the level of certain sRNAs (e.g. 6S RNA) that may act as important global regulators of transcription and play a critical role in bacterial stress responses (50,51). YbcL in uropathogenic *E. coli* has been shown to play an important role in suppressing innate immune response during urinary tract infection (52,53), and it is involved in the regulation of pathogenicity and metabolic adaptation to host conditions (54–56). CarO (Synonym: YgiW) in *P. aeruginosa* may regulate intracellular Ca2+ homeostasis, surface-associated motility, resistance to tobramycin, and the production of the virulence factor pyocynin (57). In *E. coli*, YgiW is required for cell survival in hydrgogen peroxide and cadmium (58), and it plays a role in antimicrobial resistance and virulence in *Salmonella enterica* Typhimurium (59). PgpA in *P. aeruginosa* is required for tobramycin resistance (60). Taken together, for the genes highly enriched in clade1, their hypothetical roles in pathogenicity have been verified by many previous experimental investigations. In addition, as revealed by literature, most of these genes are prevalent in several bacterial pathogens, suggesting their functional role in bacterial pathogenesis are not limited in *Pseudomonas* species.

### 4. Homogeneity analyses unveil structurally conserved genes with diversified functions

From the perspective of pan-genome analysis, both core genome and accessary genome are important genetic elements implicated in driving evolution and diversity. A large portion of accessary genome in *Pseudomonas* pan-genome analysis indicates gain-of-function and lossof-function play a role in driving genome diversity. Do core genes also play a role in driving diversity? It has been reported that conserved orthologous genes may indeed show functional diversity to certain extent (61,62). It would be informative to investigate what kind of conserved genes in *Pseudomonas* also play an important role in driving diversity.

Therefore, we performed a pan-genome based functional homogeneity analysis to detect genes that are structurally conserved in most genomes but with diversified functions, indicated by a low functional homogeneity index (see methods part for details). Considering the limitation of a small core-genome of the 704-genome data set, we started the analysis with the dataset of the 20 representative genomes defined for the comparative functional enrichment analysis (Table S5). In addition to functional homogeneity analysis, we have also calculated geometric homogeneity index of the genes, which indicates the extent of gene structural conservation. For this 20 representative genomes dataset, we only considered the single-copy core genes as structurally conserved genes. Figure 4 shows the pan-genome of the 20 representative genomes, with each genome visualized by one layer of circle. In the pan-genome, the clustering of the genome layers is based on presence/absence of orthologous gene family using Euclidean distance and Ward Linkage. Functional homogeneity index is visualized in one of the extra layers in dark green (layer name: Func. Homeogeneity Ind.). The value of functional homogeneity index for single-copy core genes varied between 0.642 and 0.999 (mean, 0.877). Geometric homogeneity index is visualized in the same way (layer name: Geo. Homeogeneity Ind.), and the value of geometric homogeneity index for single-copy core genes varied between 0.704 and 1.000 (mean, 0.965), indicating generally well-conserved gene structures of these single-copy core genes (note: the lowest value for all the gens in the pan-genome is 0.501). We have discovered only seven genes that are structurally conserved across all the genomes but show functional homogeneity < 0.7.

**Fig. 4.**
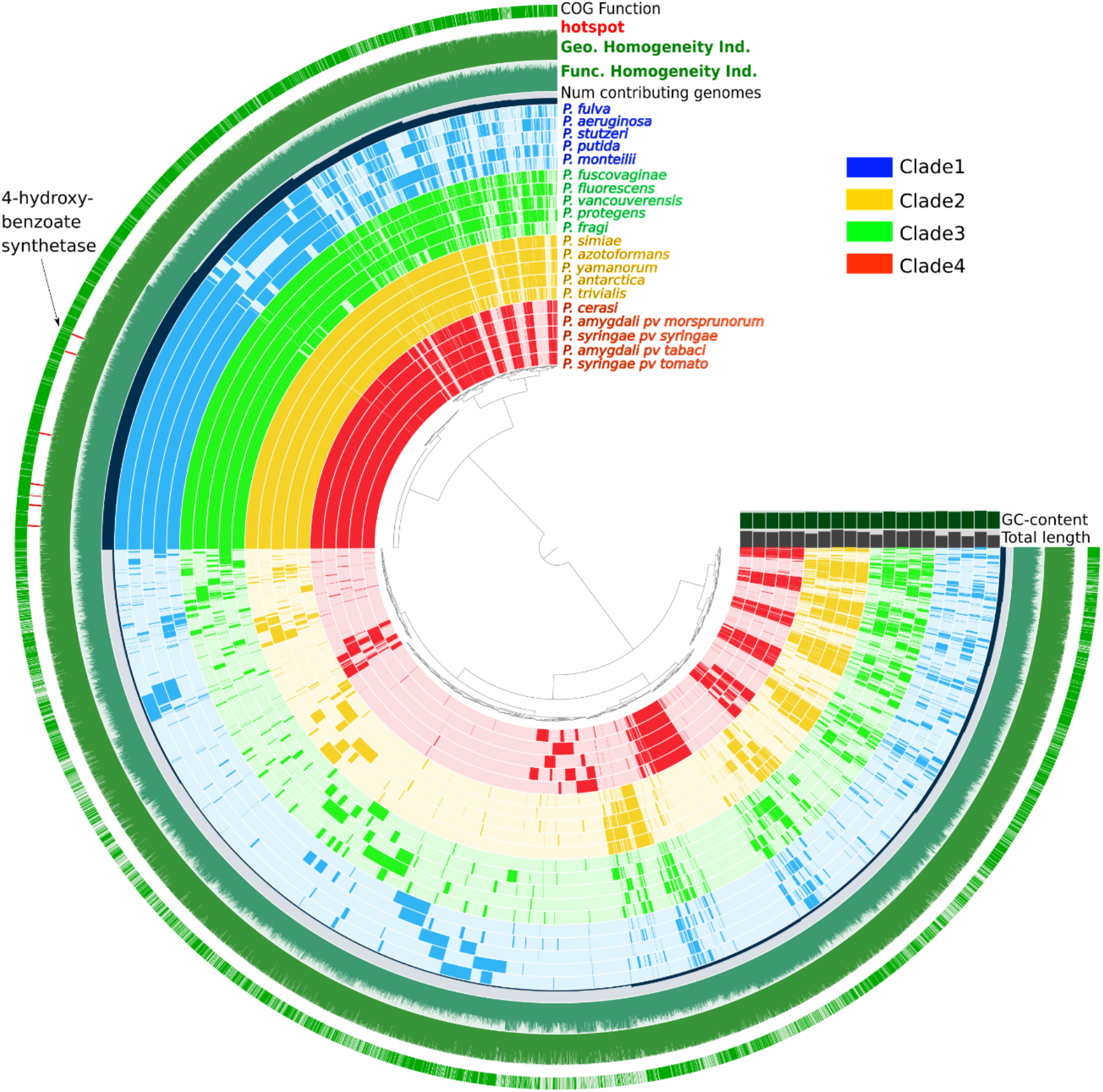
Pan-genome of 20 representative *Pseudomonas* genomes. One layer represents one genome, with a few extra layers showing distinct pan-genome features. The color of each genome layer (and layer label) is indicative of the ANI based clade that the respective species fall in. Num of contributing genomes, the number of genomes each orthologous gene family identified from (range 1-20). Func. Homeogeneity Ind., functional homeogeneity index. Geo. Homogeneity Ind., geometric homogeneity index. The layer “hotspot” indicates the exact positions of structurally conserved genes with diversified functions, and one representative gene encoding 4-hydroxybenzoate synthetase is labelled.

These genes encode proteins including 4-hydroxybenzoate synthetase (gene: ubiC), Molybdopterin-guanine dinucleotide biosynthesis protein A (gene: mobA), D-hexose-6-phosphate mutarotase (gene: yeaD_1), sulfur transfer complex TusBCD TusB component (dsrH), Flagellar hook-length control protein FliK (gene: fliK), VRR-NUC domain-containing protein (gene: HWN47_21460) and DNA uptake protein ComEA (gene: comEA) (details are listed in Table S10). Here gene names refer to representative homolog name in *P. aeruginosa*.

The exact positions of these genes are marked in the layer of “hotspot”, with one representative gene encoding 4-hydroxybenzoate synthetase labelled in the pan-genome (Figure 4). We have further verified the results with the complete dataset of the 704 *Pseudomonas* genomes. All the genes are to certain extent structurally conserved, and present in at least 300 genomes. In particular, ubiC, HWN47_21460 and mobA are structurally well conserved. They are present in 697 genomes, 691 genomes and 689 genomes, respectively.

Thus, by analyzing functional homogeneity of core genes in this study, we could detect multiple genes that show diversified functions while still keeping a relatively conserved structure across different genomes. Among these proteins, 4-hydroxybenzoate synthetase, Molybdopterin-guanine dinucleotide biosynthesis protein A, D-hexose-6-phosphate mutarotase and sulfur transfer complex TusBCD TusB component are involved in resource usage with broad substrate specificity (63–65), and Flagellar hook-length control protein FliK is implicated in regulating bacterial motility (66,67). These functions typically play an important role in environmental adaptation of the bacteria. This is consistent with the assumption that these genes are also important genetic elements driving diversity. Spontaneous mutant in these genes might be able to help the bacterium get adapted to new ecological niches, such as by achieving the ability of using new nutrient elements or moving to a new environment to get a better chance of propagation.

## Conclusion

In this study, we have analyzed the pan-genome characteristics of 704 complete *Pseudomonas* genomes, and investigated intrageneric structure of the genus *Pseudomonas* based on complete whole-genome sequence information. Further comparative functional enrichment analysis has revealed multiple functions that are predominantly enriched in opportunistic human pathogens, while largely lacking in species only living in natural habitats, suggesting the possession of these functions might be the special genomic feature of *Pseudomonas* human pathogens.

Pan-genome analysis of the 704 *Pseudomonas* genomes has revealed a small core genome (representing 1.62% of the pan-genome) and a very large accessory genome (98.38% of the pan-genome), indicating a high level of genomic diversity. Intrageneric structure analysis based on whole-genome similarity index ANI grouped the *Pseudomonas* species in four clades, which was reinforced by *Pseudomonas* phylogeny established with concatenated alignment of singlecopy core genes and by relatedness of orthologous genes between 20 representative strains, as displayed in Figure 1, 2&4. Most known opportunistic human pathogens, such as *P. aeruginosa*, *P. stutzeri and P. putida*, fall into one clade.

Moreover, in this study, we have investigated host-related functional elements. Although many *Pseudomonas* species may live in tremendously diversified environments, it is still poorly understood why some *Pseudomonas* spp. may infect humans or use animals as host, while other species only live in natural habitats. With a combination of intrageneric structure analysis and comparative functional enrichment analysis, we have identified functions that are predominantly present in human (or animal) pathogen species, while largely lacking in species only living in natural habitats. Most of these functions have been shown by previous experimental investigations to play an important role in bacterial pathogenesis, suggesting the possession of these functions might be characteristic of those *Pseudomonas* species with potential of becoming human pathogen. It is noteworthy that these functions are not only present in *Pseudomonas* pathogens, also prevalent in several other bacterial pathogens, which indicates these functions can be potential candidates for the development of new antibiotics.

In summary, the information we achieved from this study added to our understanding about the genome varieties that generate the tremendous diversity in *Pseudomonas*, and provided insights into host-related functional determinants, which might be beneficial for the development of more targeted antibiotics.

## Author contributions

BY and AD conceived and designed this study. BY performed the analyses. BY and AD wrote the paper together.

## Acknowledgements

The authors acknowledge the constructive feedback from Dr. Sébastien Boutin.

## Data Availability Statement

The datasets generated and analyzed during the current study are available in the RefSeq database of the National Center for Biotechnology Information (NCBI). Detailed information including assembly accession is listed in Supplementary Table S1.

---

**Fig. S1.**
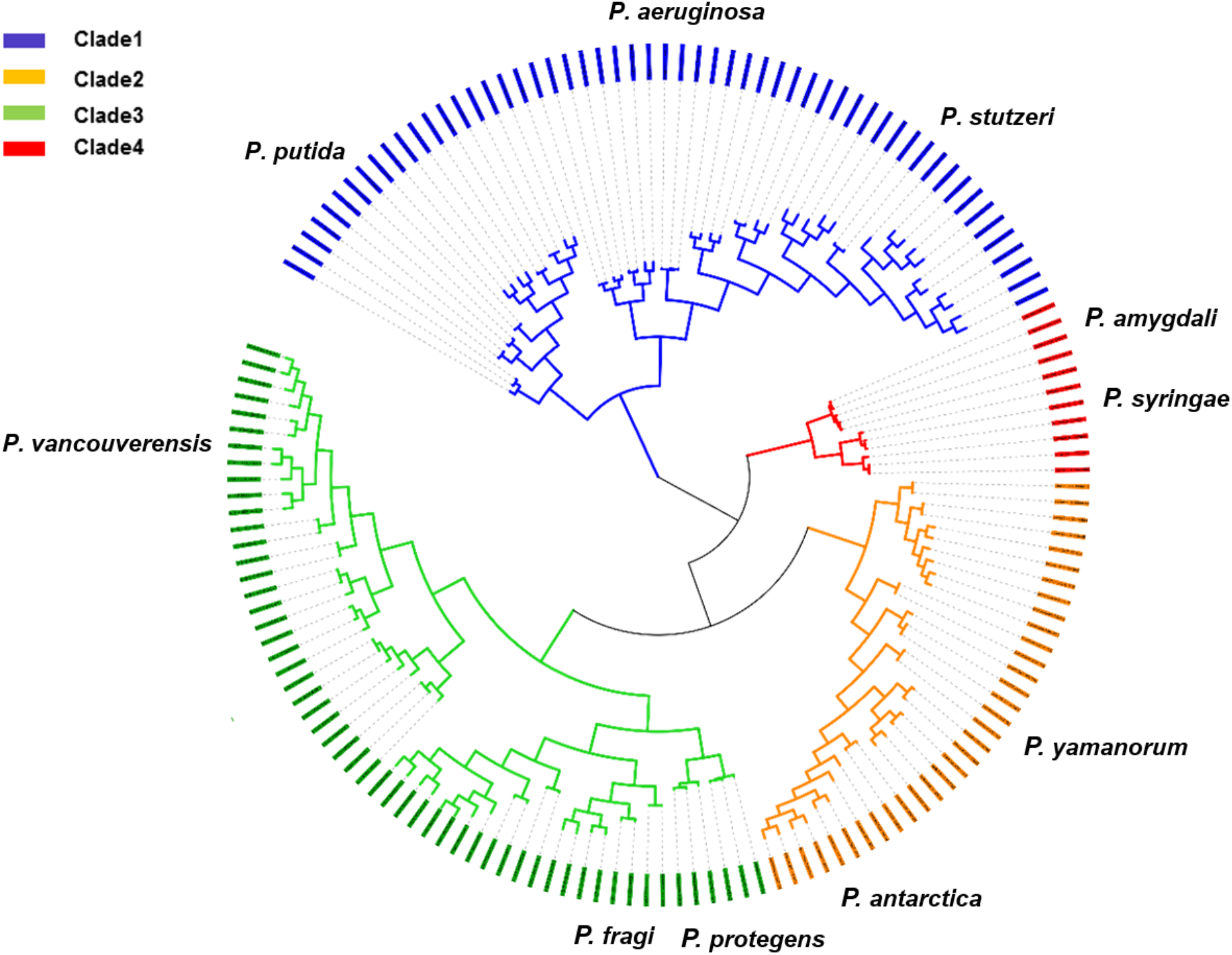
Population structure of the genus *Pseudomonas* based on whole-genome comparisons (ANIm) of 155 *Pseudomonas* strains representing 95 valid *Pseudomonas* species. The calculations were performed using the Python module PYANI. The strains fall into four major Clades (Clade1-Clade4). The positions of representative species in each Clade were labelled.

**Fig. S2.**
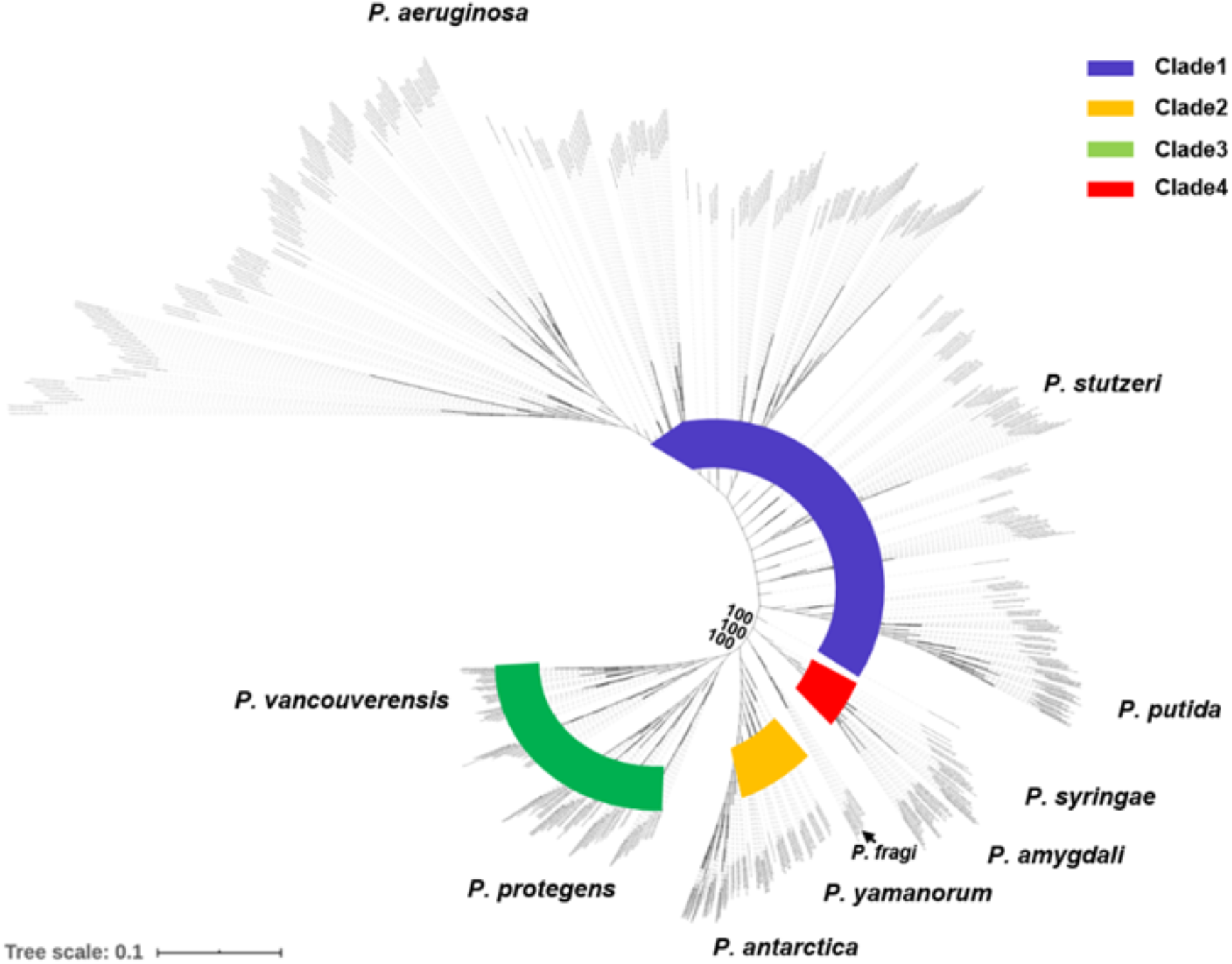
Phylogeny of 704 *Pseudomonas* strains. The phylogenetic analysis was performed using the Maximum Likelihood method based on concatenated single-copy core genes. Four Clades comparable to the four major Clades formed by whole-genome comparisons (ANI) have been detected and color-labelled. The positions of representative species in each Clade were labelled. The numbers indicate bootstrap values of the main branches.

